# An H3K79 Methylation-Dependent Checkpoint Blocks Holliday Junction Resolution and Meiotic Divisions under Heat Stress

**DOI:** 10.64898/2025.12.09.692842

**Authors:** Noopur Joshi, Neeraj Joshi, G. Valentin Börner

## Abstract

Meiosis, the specialized cell division that produces haploid gametes from diploid precursors, is markedly more sensitive to heat stress than mitosis in a wide range of organisms, despite the two processes utilizing largely overlapping cellular machinery. The mechanistic basis of meiotic heat sensitivity has remained unclear. Here, we show that in budding yeast, even moderate heat stress impairs the processing of programmed double-strand breaks into crossovers, resulting in permanent meiotic division arrest. This arrest depends on methylation of histone H3 at lysine 79 (H3K79) by Dot1, and a subset of other components of the meiotic recombination checkpoint. Thus, in contrast to the canonical heat shock response that adapts mitotically dividing cells to elevated temperatures, meiotic arrest is triggered by stalled recombination intermediates in the context of the H3K79 chromatin modification. These findings suggest a conserved epigenetic pathway responsible for heat-induced meiotic failure across eukaryotes. Genetically enhancing the heat tolerance of meiotic chromosome metabolism may increase reproductive resilience of organisms facing rising global temperatures, bridging the gap until mitigation strategies are in place.

**Significance Statement:** Meiosis generates haploid gametes through a specialized cell division that involves crossovers between homologous chromosomes. Crossovers promote genetic diversity and ensure proper chromosome segregation during meiosis I. In many organisms, meiosis stalls under moderate heat stress at temperatures permissive for mitosis, a phenomenon with implications for crop fertility and stability of ecosystems. Using yeast, we discovered that heat stress blocks meiosis at Holliday junctions - the four-armed DNA intermediates that resolve into crossovers - while alternative non-crossovers form normally. Arrest depends on the H3K79 methylation-dependent checkpoint involving Dot1. Thus, heat sensitivity of meiosis stems from checkpoint surveillance of impaired chromosome interactions. Modulating the underlying mechanism could enhance reproductive resilience to a warming climate.

## Introduction

Meiosis is the specialized cell division that reduces diploid chromosome content to haploid across eukaryotes, as a prerequisite for sexual reproduction (1). During prophase I, recombination generates crossovers between homologous chromosomes (homologues) which are critical for their separation to opposite spindle poles during the first meiotic division (MI). Crossovers arise from the repair of programmed DNA double-strand breaks (DSBs), introduced by the conserved Spo11 protein complex (2). Only a subset of DSBs is processed into crossovers, via double Holliday junction intermediates whereas other DSBs are processed via less stable strand exchange events into non-crossovers, i.e. inter-homologue gene conversions without exchange of flanking chromosome arms (3, 4). While the homologue is preferred as recombination partner during meiosis, the sister chromatid may serve as repair template during early wild-type meiosis and under mutant conditions, via a recombination pathway that also involves double Holliday junctions (5, 6).

The MI cell division is closely coordinated with the orderly progression of recombination, as part of a broader program of chromosome morphogenesis that also includes homologue pairing and synapsis (1). Mutants defective at intermediate steps of DSB processing are either transiently or permanently arrested during prophase I, a block mediated by the meiotic recombination checkpoint, sometimes referred to as pachytene checkpoint (7–10). This meiotic arrest and/or delay can be triggered by diverse defects in chromosome metabolism, ranging from unresected (11) or hyper-resected DSBs (8) to stalled later recombination intermediates (12). These defects result in persistent Mec1^ATR^ and/or Tel1^ATM^-mediated phosphorylation of meiotic chromosome axis protein Hop1 which in turn recruits and activates CHK2-related kinase Mek1 (13). Activated Mek1 then phosphorylates Ndt80, blocking its transcriptional activity required for prophase I exit (14).

Many meiotic recombination and checkpoint components are shared with those active in the DNA damage response in mitotically proliferating cells (8, 15). Checkpoint mutants encompass factors that monitor recombination progression, but they may also mediate pathway choice for DSB processing, blurring the distinction between pure surveillance factors and mechanisms that control repair of the underlying lesion, with secondary effects on meiotic progression (6, 12, 16, 17). Importantly, because checkpoint effects have been examined mainly under mutant conditions, the degree to which they function during normal wild-type meiosis remains an open question (10, 12).

Meiosis is unusually sensitive to even moderate heat stress across fungi, animals, and plants (18–24), unlike mitosis which at the same temperatures proceeds efficiently in the same organism (25–28). Accordingly in budding yeast, mitosis can occur up to 42°C, whereas meiotic divisions are blocked around 35°C (21, 25). Similar patterns have been observed in multicellular organisms: exposure of the crop plant wheat to 30°C specifically during early meiosis results in progression block and grain loss (18), and in the nematode *C. elegans* at 26°C, synaptonemal complex assembly and homologue linkage via crossovers are disrupted (24), even though mitotic divisions in both organisms remain intact at the respective temperatures (26, 28). This pattern suggests that a conserved feature of the meiotic core program is responsible for the heat sensitivity of meiosis rather than organism-specific vulnerabilities.

Whether heat tolerance of meiosis can be genetically modulated and what mechanisms underlie this trait, remains poorly understood. In this study, we show that rather than responding to cytoplasmic protein denaturation, heat-induced meiotic arrest is a chromatin-based checkpoint that shares features with the meiotic recombination checkpoint. We uncover a central role for the epigenetic mark methylated histone H3 at lysine 79 (H3K79me), deposited by Dot1 (15, 29–31), in both meiotic arrest and germ cell survival under moderate heat stress. Thus, survival of wild-type germ cells during environmental heat stress is secured by a checkpoint previously studied only in mutation-induced conditions (10). We further establish that moderate heat stress attenuates two key meiotic recombination steps, the formation of DSBs and the resolution of double Holliday junctions into crossovers, and that Dot1 is responsible for this attenuation, an effect separable from its checkpoint function. In summary, our study identifies a novel, meiosis-specific heat stress response pathway that is both spatially and mechanistically distinct from the canonical mitotic heat shock response (32). This work provides a framework for understanding how epigenetic signaling governs reproductive heat resilience.

## Results

### Recombination-associated events trigger meiotic arrest during moderate heat stress

Effects of heat stress on meiosis were investigated in the budding yeast *S. cerevisiae*, a model organism where meiotic divisions and recombination can be analyzed in the same cell population (4). Recognizing that the temperature above which meiotic divisions fail may vary among different species and strains, we began by determining the lowest restrictive temperature for the SK1 wild-type strain used in the current study. When diploid cultures arrested at the G1 stage are transferred to meiosis medium containing a non-fermentable carbon source, cells undergo efficient and synchronous meiosis (33). At 30°C, the reference temperature used in this and other studies, ≥ 80% of cells have undergone one or both meiotic divisions by 8.5 h as indicated by high abundance of cells containing 2 or 4 nuclei, and typically ∼90% of cells have completed meiosis by t = 12 h (Fig. 1A to C). To allow premeiotic replication at 30°C, a master culture was incubated in meiosis medium until t = 2 h (34), followed by a shift of aliquots to closely spaced, elevated temperatures above 33°C, a temperature at which the SK1 strain readily completes meiosis (4, 17) (Fig. 1B). At temperatures up to 34°C, most cells completed meiosis, with a 0.8°C increase above 33.2°C reducing meiotic divisions by less than 30% (Fig. 1C). However, a further 0.8°C increase from 34.0°C to 34.8°C led to an approximately 10-fold reduction of meiotic divisions, with cells remaining arrested even after prolonged incubation (Fig. 1C; Fig. S1A). Thus, wild-type meiosis exhibits a sharp temperature threshold for MI divisions at ∼34.8°C, which henceforth was used as the lowest restrictive temperature.

**Fig. 1.**
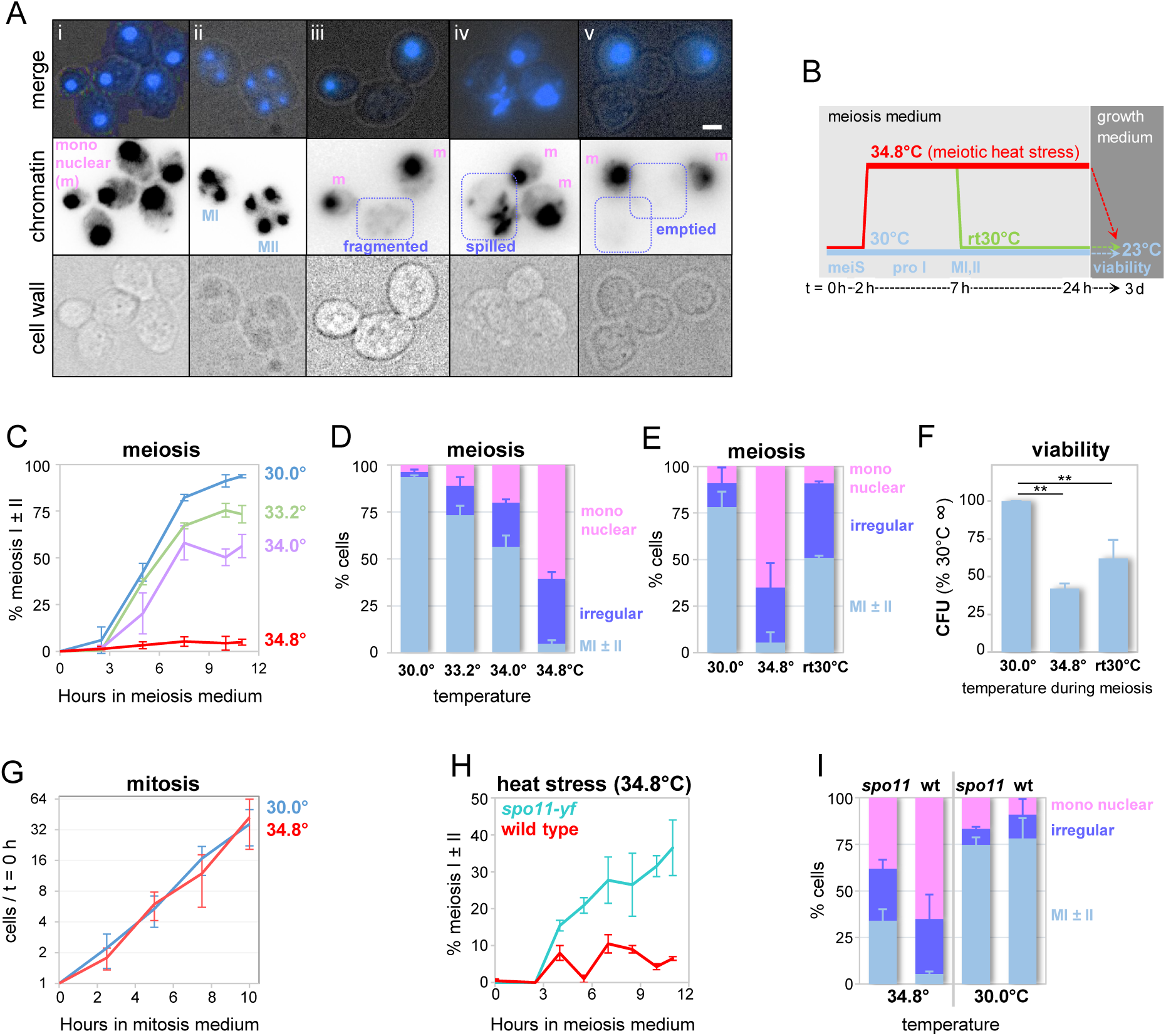
Moderate heat stress triggers meiosis-specific viability loss and division arrest in response to programmed DSBs. (A) Representative images of two to six wild-type cells carrying **(i)** one (mononuclear), **(ii)** two (MI) or four nuclei (MII) as well as irregular nuclear morphologies exhibiting **(iii)** fragmented chromatin, **(iv)** burst cells with protruding chromatin (“spilled”) and cells apparently lacking chromatin (“emptied”). Chromatin and cell wall status were assessed using epifluorescence and brightfield microscopy, respectively. Size bar is 1 μm. (B) Diagram for incubation protocols. See text for details. (C) Efficiency and kinetics of meiotic divisions as indicated by cells with two or four nuclei in synchronized cultures at temperatures ranging from 30°C to 34.8°C. Cultures are derived from a single starter culture. n = 3; error bars are SDs. (D) Percentage of cells at late time points exhibiting regular [mono-nucleate or divided (two or four nuclei)] or irregular morphologies (see Fig. 1A). Cultures are the same as in Fig. 1C, averages of t = 10 h and 11 h are shown (n = 3; error bars are SDs. For kinetics of irregular morphologies and individual irregular classes see **Fig. S1B,C** respectively). (E) Partial reversibility of the heat-induced meiotic arrest. Aliquots from parallel meiotic cultures were incubated at 30°C, at 34.8°C, or shifted at t = 7 h from 34.8° to 30.0°C (rt 30°C). Percentages of regular or irregular cell morphologies are averages of t = 10 h and 11 h, except for the shifted culture which reached final levels at t = 11 h. n = 2, error bars indicate range. (F) Permanent and transient heat exposure during meiosis result in viability loss. X-axis indicates incubation temperatures during meiosis. Colony-forming units (CFU) were determined following transfer to solid growth medium and incubation at 23.0°C. Values are expressed as percentages relative to the control culture undergoing meiosis at 30°C [3.2 (± 0.1) x 10^7^ cfu/ ml). Asterisks indicate p-values <0.01 as determined by one-tailed paired Student’s t-test. For timing of meiotic divisions see **Fig. S1D**. (G) Cell numbers in mitotic cultures normalized to t = 0 h incubated in liquid growth medium (YPA) at 30°C and 34.8°C. Average cell concentration was 2.5 (± 0.9) x 10^5^ /ml at t = 0 h. n = 2, error bars indicate range. (H) Meiotic arrest under heat stress is triggered in part by event(s) associated with meiosis-specific DSBs. Graph indicates percentage of cells that have completed one or both meiotic divisions in cultures carrying wild-type or catalytically dead *SPO11 (spo11-yf*) (n = 2; error bars indicate range). See Fig. **S1F** for meiotic divisions at 30°C of subcultures derived from the corresponding master cultures. A transient dip from 10% to 1% observed in some WT cultures (here at t = 5.5h) correlates with a peak in irregular cells (Fig. S1E), likely due to morphological similarities between binucleate and fragmented cells. (I) Percentage of *spo11-yf* and wild-type cells analyzed in Fig. 1H exhibiting regular or irregular cell morphologies at late time points (averages of t = 10 h and 11 h; n = 2; error bars indicate range). See **Fig. S3B-v** for irregular morphology classes.

Following incubation at 34.8°C until t = 24 h, most cells remain mononucleate, consistent with prophase I arrest (21), yet about one third of bona-fide undivided cells exhibit one of several irregular morphologies, including fragmented chromatin (“fragmented”), chromatin attached to a burst open cell (“spilled”), or cell ghosts lacking chromatin (“emptied”; Fig. 1A,iii to v; Fig. S1B,C). Such irregular cells are rare at 30°C but at 34.8°C have reached substantial levels by t = 5 h (Fig. 1D; Fig. S1B,C). Moreover, incubation at the lowest restrictive temperature until t = 24 h followed by transfer to mitotic growth medium results in a dramatic loss in viability, as evidenced by a ≥ 2-fold decrease in colony formation following meiosis at 34.8°C compared to the viability in cognate reference cultures (“30.0°C”; Fig. 1B,E,F). Conversely, parallel G1-arrested cultures transferred to fresh growth medium instead of meiosis medium at t = 0 h, undergo mitotic cell divisions with essentially identical kinetics at 34.8°C and 30°C, indicating comparable viabilities (Fig. 1G). These findings are qualitatively similar to earlier reports analyzing a different yeast strain background showing progressive loss of spore viability with increasing duration of heat exposure during meiosis (35).

We conclude that that wild-type meiosis exhibits a sharp thermal threshold under relatively moderate heat stress, with meiotic efficiency dropping from near-complete divisions to permanent arrest over a 1.6°C range. By contrast, the G1 arrest triggered by the canonical heat shock response at temperatures above 37°C is transient (36), and mitotic rates decline progressively between 37°C and 45°C (25).

Two lines of evidence suggest that meiotic divisions are blocked as part of a regulatory cell division response, rather than being caused by an inherent heat sensitivity of the meiotic machinery: First, most cells complete meiosis when returned to permissive conditions following heat stress for five hours (“rt30°C”; Fig. 1B,E; Fig. S1D) (21). Second, elimination of recombination via a catalytically inactive *SPO11* allele (*spo11-yf*) (37) partially restores nuclear divisions at 34.8°C (Fig. 1H). Notably, in wild-type rt30°C and in *spo11-yf* cultures at 34.8°C, the arrest is bypassed only in a subset of cells, whereas the percentage of cells exhibiting irregular morphologies remains unchanged under both conditions, suggesting that progression is restored in regular mononucleate cells only (Fig. 1E,I; Fig. S1E).

We conclude that the meiotic heat stress response entails two distinct processes: First, defects in Spo11-induced meiotic recombination and/or associated events trigger a reversible cell division arrest at the mononucleate stage. Second, some cells undergo irreversible morphological changes independent of recombination. Subsequent analyses focused on the recombination-dependent effects.

### Heat stress attenuates DSB formation and crossover-specific double Holliday junction resolution

Given the role of DSB-initiated events in triggering meiotic arrest during moderate heat stress, recombination was monitored in the same cultures analyzed for meiotic divisions (Fig. 2; Fig. S1D). Physical analyses of recombination intermediates and products at the well-characterized *HIS4LEU2* DSB hotspot (Fig. S2A) (38, 39) revealed that heat stress dramatically affects three transitions in recombination: First, DSBs are decreased 2.5 (± 0.7)-fold at 34.8°C compared to 30°C in *sae2*Δ, a mutant background that accumulates DSBs due to defective resection and ensuing repair (7, 40) (Fig. 2A,B). Following the shift to 34.8°C, DSB formation stops almost immediately, while continuing for three more hours in cognate reference cultures at 30°C (Fig. 2B). The DSB reduction under moderate heat stress is unlikely to result from defective pre-meiotic replication, which appears to be mostly complete at the time of shift to 34.8°C at t = 2 h. Accordingly, in *SAE2* reference cultures at 30°C, DSB steady-state levels reach their peak by t =2.5 h (Fig. 2C,D-i), and DSBs form only after premeiotic replication is complete (41). To account for the ∼2.5-fold decrease in DSB levels, y-axes for post-DSB intermediates and products at 34.8 °C were proportionally expanded relative to those at 30 °C (Fig. 2D; for unadjusted axes see Fig. S2C).

**Fig. 2.**
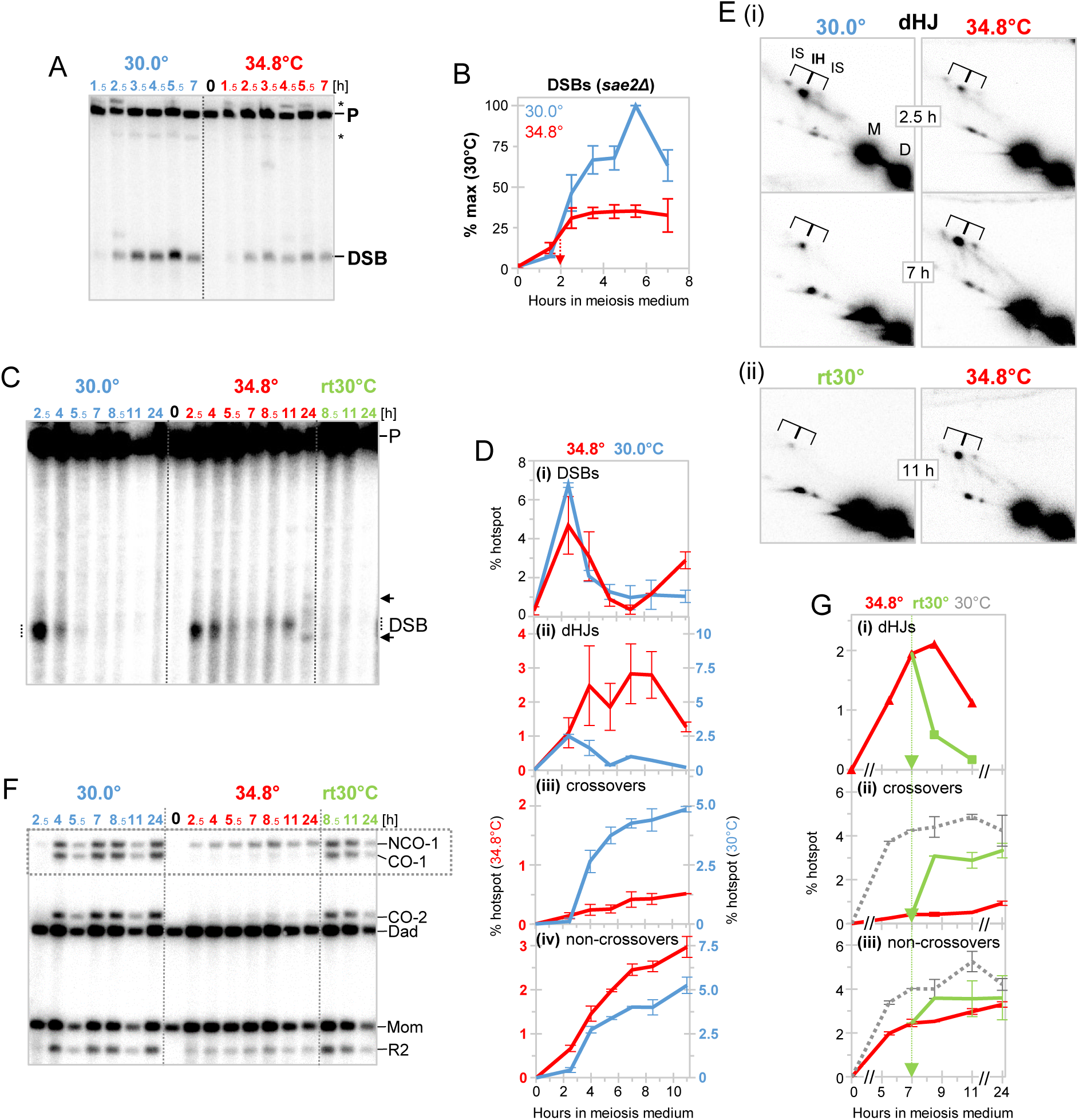
Heat stress attenuates DSB formation and Holliday junction resolution into crossovers. (A) Southern blot analysis of DSB accumulation in *sae2*Δ at 30°C and 34.8°C at the *HIS4LEU2* hotspot in PvuII-digested genomic DNA. For a diagram of the *HIS4LEU2* hotspot see **Fig. S2A**. A meiotic culture was split after taking the t = 0 h sample (middle) and subcultures were incubated under the indicated conditions. P, parental. Asterisks indicate cross-hybridizing bands. (B) Quantitative analysis of DSBs in *sae2*Δ reveals rapid cessation of DSB formation upon heat exposure. DSBs are shown as percent of the maximum levels in the corresponding meiotic culture incubated at 30°C [24.5 (± 6.6)% of DNA; n = 3; error bars indicate SD]. DSB levels at 34.8°C compared to 30.0°C are 2.5 (± 0.7)-fold reduced at times when DSBs have reached their respective plateau levels [at 4.5 h (34.8°C) and 5.5 h (30.0°C), respectively; n = 7]. Red arrow indicates the time of culture shift from 30° to 34.8°C. (C) Southern blot analysis of DSB steady state levels in split wild-type cultures (*SAE2*) incubated at 30°C, 34.8°C, or shifted from 34.8°C to 30°C at t = 7 h (rt30°C). (D) Quantitative analysis of recombination intermediates and products at the *HIS4LEU2* hotspot. Higher DSB levels at 30°C compared to 34.8°C (see Fig. 2B) are accounted for by expanding the y-axis for 30°C 2.5-fold (right) for post-DSB intermediates and products. For unadjusted y-axes, see **Fig. S2C**). (i) DSB steady state levels quantitated from PvuII-digested genomic DNA (see Fig. 2C); (ii) total (interhomolog and intersister) double Holliday junctions, as quantitated from XhoI Southern blots. For representative excerpts see Fig. 2E and for complete blots **Fig. S2B**, (iii) crossovers, (iv) non-crossovers (CO-1 and NCO-1, respectively, in Fig. 2F). n = 2, error bars indicate ranges. (E) Two-dimensional gel Southern blot analysis at *HIS4LEU2* of XhoI-digested genomic DNA at informative time points. **(i)** Cognate WT cultures at 30°C and 34.8°C; (ii) cultures shifted to 30°C at t = 7 h (rt30°, left) or kept at 34.8°C (right) at t = 11 h. Interhomolog (IH) and intersister (IS) double Holliday junction positions are indicated. For complete blots see **Fig. S2B**. (F) One-dimensional gel Southern blot analysis of crossovers and noncrossovers at *HIS4LEU2* in genomic DNA digested with XhoI and BamHI in meiotic subcultures incubated at 30°, 34.8°C, as well as following a shift at t = 7 h from 34.8° to 30°C (rt30°C). The dotted box indicates representative crossover (CO-1) and non-crossover bands (NCO-1). (G) Quantitative analyses of effects of a transfer of cultures from 34.8° to 30°C on **(i)** double Holliday junctions, **(ii)** crossovers, and **(iii)** non-crossovers. Green arrows indicate the time of shift to 30°C (t = 7 h). The gray dotted curves in (ii) and (iii) indicate COs and NCOs in the cognate reference cultures at 30°C. X-axes are interrupted before 5 h and after 11 h.

Second, DSB strand exchange at 34.8°C is impaired. This is indicated by persistence of a small fraction of DSBs at late time points in WT (*SAE2*; Fig. 2C,D-i), a more moderate reduction in DSB peak levels in this background than expected from analysis in *sae2*Δ, consistent with transient DSB accumulation, and by the delayed peak at 34.8°C versus 30°C of total double Holliday junctions [dHJs; comprising dHJs formed between homologues (IH) and those between sister chromatids (IS)] (Fig. 2E-i, 2 D-ii) (39).

Third, and importantly, dHJs persist under heat stress at substantially higher levels compared to 30°C (Fig. 2D-ii). Moreover, crossovers are essentially eliminated under heat stress, whereas remarkably, non-crossovers form with comparable efficiencies as at 30°C (Fig. 2F; 2D-iii,iv). Upon shift from 34.8°C to permissive conditions, persisting dHJs are rapidly resolved (rt30°C; Fig. 2E-ii, 2G-i; Fig. S2B), and crossovers are restored to 70% of the cognate 30°C culture, with negligible effects on non-crossovers (Fig. 2F, G-ii,iii).

We conclude that DSB formation and DSB processing along the crossover pathway, including dHJs resolution into crossovers, are attenuated under moderate heat stress, whereas DSB processing into non-crossovers is largely unaffected. These findings further demonstrate that dHJs predominantly generate crossovers during WT meiosis whereas non-crossovers are formed via a dHJ-independent pathway, a model previously inferred from studies in mutant meiosis (3, 4).

### Recombination status during heat-stress is monitored via Dot1-mediated H3K79 methylation

We next sought to identify factors other than *SPO11* that contribute to the heat-stress-induced arrest during wild-type meiosis. To determine whether this surveillance pathway overlaps with the checkpoint activated by mutation-induced recombination defects, we performed a candidate screen targeting known or suspected components of the meiotic recombination checkpoint, including the chromosome axis modulator Pch2^TRIP13^ (9, 12, 42) and the histone methyltransferase Dot1 (10, 29). In cell patches on solid sporulation medium at 35°C, UV fluorescence of spore walls [a visual marker of meiotic progression (4)] was weak in WT and absent in *pch2*Δ, but detectable in *dot1*Δ, even though fluorescence levels under permissive conditions were indistinguishable for the three genotypes (18°C; Fig. 3A). Importantly, meiotic divisions in *dot1*Δ at 34.8°C were increased 4- to 5-fold compared to WT (Fig. 3B-i), whereas no differences were observed in the cognate subcultures at 30°C (Fig. S3A-i). While mononucleate cells in *dot1*Δ were substantially decreased compared to WT, cells with irregular morphologies were essentially identical between the two genotypes, suggesting that *DOT1* mediates arrest under heat stress in mononucleate cells only (Fig. S3C). Unlike in *dot1*Δ and WT, meiosis in *pch2*Δ is partially arrested even at the highest permissive temperature of 33°C, an arrest response triggered by defects in *SPO11*-mediated recombination events (6, 12).

**Fig. 3.**
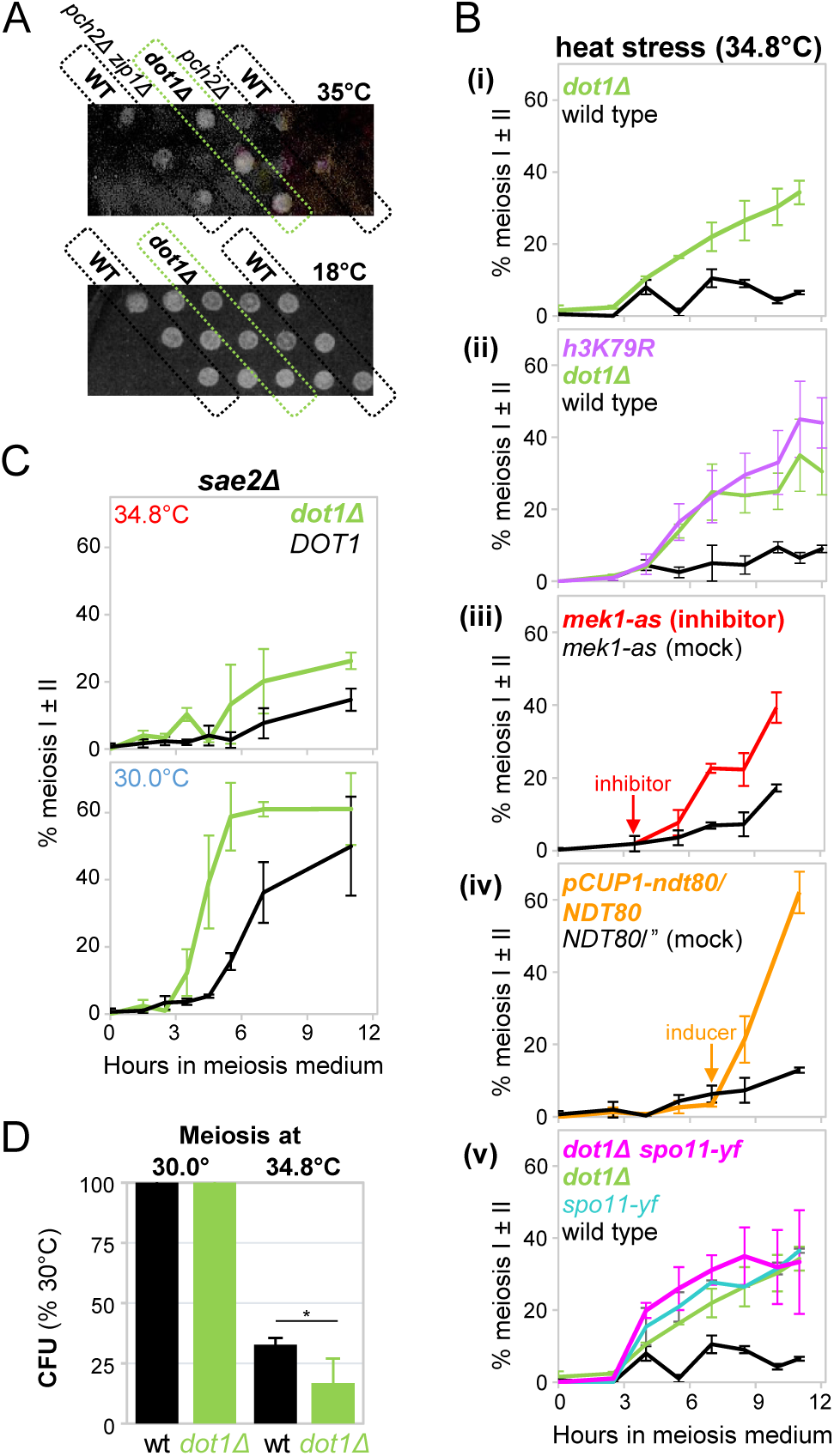
The heat-induced checkpoint arrest depends on Dot1-mediated H3K79 methylation. (A) Increased spore wall fluorescence in *dot1*Δ under heat stress compared to wild type, but not in *pch2*Δ. Cell patches of the indicated genotypes were stamped in diagonal rows (n = 3) on rich solid sporulation medium, incubated at 35°C (top) or 18°C (bottom), and monitored for spore autofluorescence under UV light. (B) Percentages of cells in cultures of the indicated genotypes undergoing one or both meiotic nuclear divisions during heat stress (34.8°C). (i) *dot1*Δ versus WT (n = 2), (ii) *h3k79r* versus *dot1*Δ and WT (n = 2), two *h3k79r* strains (NOJ1547, NOJ1548) were analyzed each in duplicate, with essentially identical results, (iii) ATP analog-sensitive *mek1-as1* plus inhibitor added at t = 3.5 h versus *mek1-as1* mock-treated subculture (n = 3), (iv) culture heterozygous for *pCUP1-ndt80/pNDT80-NDT80* induced at t = 7 h with copper sulfate compared to an identically-treated homozygous *pNDT80-NDT80* (WT) culture (n = 3), (v) *spo11-yf dot1*Δ double mutant compared to the corresponding single mutants and wild type (n = 2). Error bars indicate ranges or SDs for n = 2 and n ≥ 3, respectively. For the cognate cultures at 30°C see **Fig. S3A**, and for irregular classes at 34.8°C see **Fig. S3B**. (C) *dot1*Δ is a classical checkpoint mutant as indicated by bypass of meiotic delay and/or arrest in resection mutant *sae2*Δ (n = 3, error bars indicate SD). (D) Absence of *DOT1* reduces cell viability following meiosis under heat stress. Viability of meiotic cells at 34.8°C (right) is indicated by the percentage of colony forming units in the cognate subcultures incubated at 30°C. Viabilities of WT and *dot1*Δ at 30.0°C were 1.7 (± 0.04) and 1.4 (± 0.54) x 10^7^ cfu/ml, respectively. p = 0.019, as determined by a paired one-tailed Student’s t-test. Error bars are SDs (n = 4).

Meiotic division arrest may be bypassed due to (i) loss of a checkpoint, (ii) modification of the defect monitored by the checkpoint, as e.g. observed in *spo11-yf*, or (iii) a combination thereof. To distinguish between these possibilities, *dot1*Δ effects on meiosis were examined in *sae2*Δ where meiotic divisions are delayed and decreased (7, 40). Remarkably, in *dot1*Δ *sae2*Δ, meiotic divisions at 30°C are substantially accelerated and enhanced compared to the *sae2*Δ single mutant (Fig. 3C-bottom). Moreover, under moderate heat stress (34.8°C), *dot1*Δ also partially bypasses arrest in *sae2*Δ (Fig. 3C-top). Notably, DSB processing is blocked under both conditions in the *sae2*Δ background (below). We conclude that *DOT1* functions as part of a classical checkpoint, as the arrest bypass observed in *dot1*Δ cells occurs without altering the underlying recombination defect.

To examine the physiological relevance of the *DOT1*-mediated checkpoint arrest, the viability of WT and *dot1*Δ cells exposed to moderate heat stress during meiosis was determined. Following meiosis at 34.8°C, cells were transferred to growth medium at t = 24 h (see Fig. 1B). Remarkably, *dot1*Δ exhibited a substantially larger loss in viability compared to the WT (Fig. 3D). Thus, by blocking meiosis under moderate heat stress, the *DOT1* checkpoint maintains viability of cells undergoing gametogenesis.

Dot1 is the only known methyltransferase responsible for mono-, di-, and tri-methylation of histone H3 at lysine (K)79, located in the nucleosome core; K79 further is the only known Dot1 substrate (29). Rendering K79 non-methylatable by replacing lysine with arginine, the *h3k79r* mutant, similar to *dot1*Δ, bypassed meiotic arrest at 34.8°C while mutants and WT were again indistinguishable at 30°C (Fig. 3B-ii; Fig. S3A-ii). Thus, Dot1 controls meiotic progression under heat stress via methylation of H3K79.

Unlike *PCH2*, several other components of the surveillance mechanism that blocks meiosis in recombination-defective mutants also contribute to the meiotic heat stress response identified here. Mek1, a meiosis-specific CHK2-related chromosomal kinase, self-activates and is recruited to chromatin in a Dot1- and H3K79 methylation-dependent manner (43, 44). Consistent with Mek1’s role in relaying the arrest signal, inhibition of an ATP-analog sensitive *mek1-as1* allele partially bypasses the meiotic arrest under heat stress (Fig. 3B-iii). Similarly, induced expression of *NDT80*, a prophase I exit factor and substrate of phosphorylation by Mek1 (14, 45), results in efficient arrest bypass (Fig. 3B-iv; Fig. S3A-iv), likely by outcompeting Mek1 kinase-mediated Ndt80 inhibition (14). Bypass of meiotic arrest by overexpression of Ndt80 was previously also observed in SC mutant *zip1*Δ (46). By contrast, Rad9^53BP1^ which binds to methylated H3K79 as part of the DNA damage response in vegetative cells (31), does not contribute to meiotic arrest in response to heat stress (Fig. S3A-vi). Finally, meiotic bypass occurs with similar efficiencies in the *dot1*Δ *spo11yf* double mutant as in the respective single mutants, revealing that the two genes work along the same pathway (Fig. 3B-v).

We conclude that moderate heat stress during meiosis attenuates recombination initiation, but also disrupts the processing of Spo11-induced recombination intermediates, activating a checkpoint response dependent on Dot1-mediated H3K79 methylation, which in turn promotes Mek1 kinase activation and inhibition of Ndt80. The checkpoint identified here represents a distinct regulatory pathway, as it operates independently of both Pch2, a key component of the meiotic recombination checkpoint, and of Rad9^53BP1^, a central mediator of the DNA damage checkpoint.

### H3K79 methyltransferase Dot1 attenuates DSB formation and double Holliday junction resolution

We next examined *DOT1* effects on recombination under standard conditions, as well as under moderate heat stress. Unexpectedly, at 30°C in a *sae2*Δ background, DSBs in *dot1*Δ accumulate to levels ∼50% above those in *DOT1* (Fig. 4A,B). In the same *dot1*Δ *sae2*Δ cultures exposed to heat stress, DSBs are similarly increased, but accumulation is largely limited to the time window prior to t = 2 h when cultures are still at 30°C, and few if any additional DSBs form after the shift to 34.8°C (Fig. 4B). Thus, *DOT1* attenuates DSB formation, while DSB formation in *dot1*Δ remains heat sensitive.

**Fig. 4.**
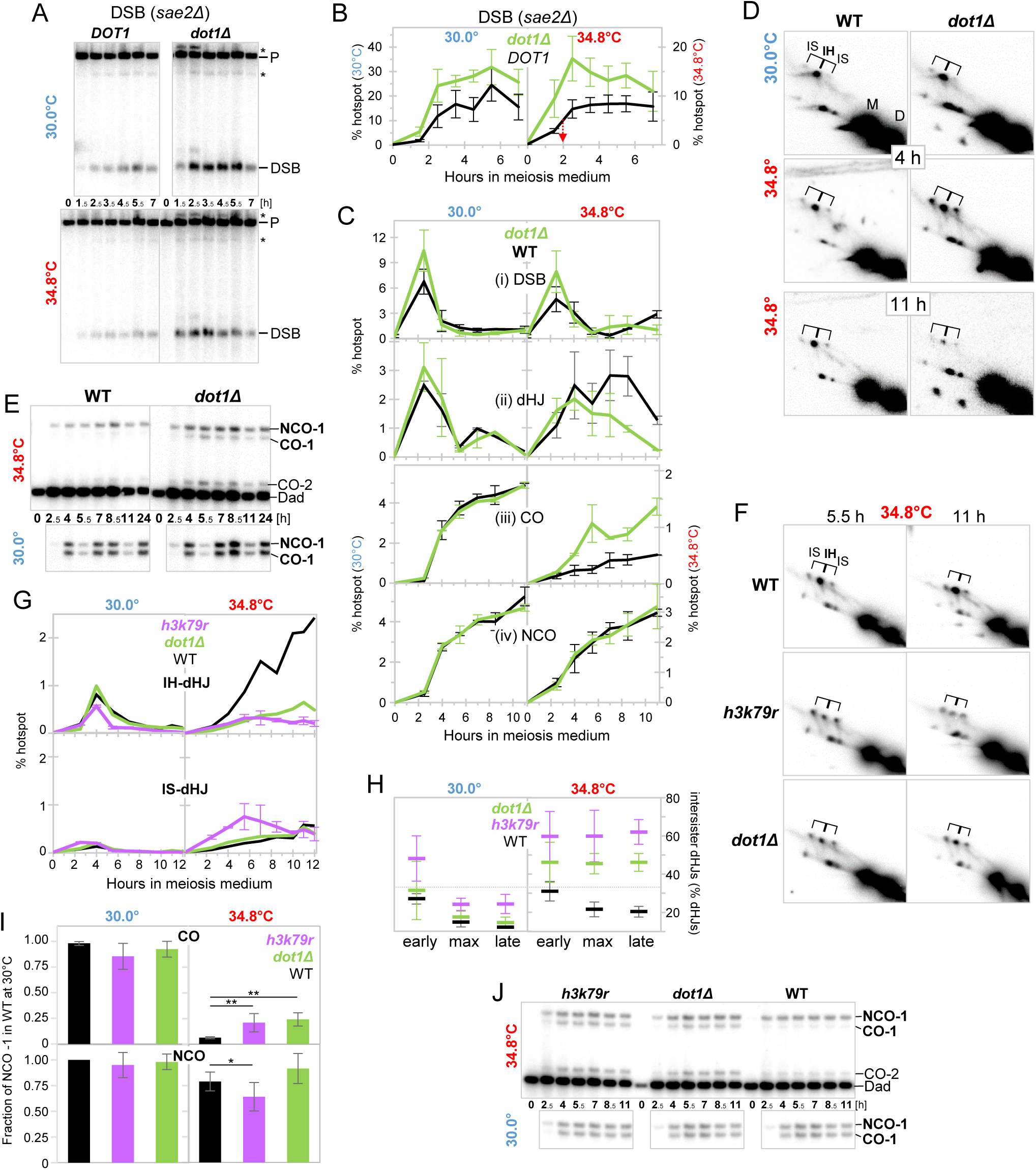
Histone H3K79 methyltransferase Dot1 attenuates DSB formation and double Holliday junction resolution into crossovers during heat stress. (A) Effect of *dot1*Δ on DSB formation in the *sae2*Δ background. 1D-gel Southern blot analyses at the *HIS4LEU2* DSB hotspot in PvuII-digested genomic DNA. After taking t = 0 h samples, both *dot1*Δ and WT cultures were split into subcultures, which were incubated at 30°C (top) or 34.8°C (bottom), respectively. P, parental; asterisks, cross-hybridizing species. (B) Quantitative analysis of DSB accumulation in *dot1*Δ and *DOT1* in the *sae2*Δ background at 30°C and 34.8°C. Red arrow indicates the time of culture shift from 30° to 34.8°C. DSB levels are provided as percent of DNA at the *HIS4LEU2* locus (n = 3; error bars indicate SD). (C) Quantitative analysis at 30°C and 34.8°C in WT and *dot1*Δ in the *SAE2* background of steady state levels of (i) DSBs, (ii) double Holliday junctions between homologues and sister chromatids; and accumulation of (iii) crossover (CO-1) and (iv) non-crossover (NCO-1) products (n = 2; error bars indicate range). (D) Two-dimensional gel Southern blot analysis of XhoI-digested genomic DNA at *HIS4LEU2* in WT and *dot1*Δ at 30°C and 34.8°C at an early (t = 4 h) and a late time point (t = 11 h). Interhomolog (IH) and intersister (IS) double Holliday junction signals are indicated. (E) 1D-gel Southern blot analysis of representative crossover and non-crossover bands in WT and *dot1*Δ at 34.8°C (top) and 30°C (bottom) following digest of genomic DNA with XhoI and BamHI. (F) Two-dimensional gel Southern blot analyses of XhoI-digested genomic DNA in WT, *h3k79r,* and *dot1*Δ at 34.8°C at early (t = 5.5 h) and late time points (t = 11 h). Interhomolog (IH) and intersister (IS) double Holliday junction signals are indicated (see **Fig. S4A f**or complete 2D blots at 34.8°C and 30.0°C). (G) Quantitative analysis of double Holliday junctions between homologues (IH) and sister chromatids (IS) at 30° and 34.8°C in *h3k79r* (n = 2, error bars indicate range), *dot1*Δ, and WT. (H) Frequency in *dot1*Δ, *h3k79r* and WT at 30°C (n = 3) and 34.8°C (n ≥ 6) of intersister double Holliday junctions as a percent of total (IS plus IH) double Holliday junctions. Averages are shown for time of first detection (early), maximum levels (max), and latest time point of reliable detection (late). Error bars are SD. (I) Effects of *h3k79r* and *dot1*Δ on maximum crossovers and non-crossovers at 30°C and 34.8°C. CO-1 and NCO-1 bands are expressed as fraction of the NCO-1 band in WT at 30°C. p-values were determined using a two-tailed t-test, with one and two asterisks indicating p ≤ 0.05 and p ≤ 0.01, respectively. (J) Informative excerpts from 1D gel Southern blot analysis of crossovers and non-crossovers in *h3k79r*, *dot1*Δ, and WT at 34.8°C (top) and 30°C (bottom) following digest of genomic DNA with XhoI and BamHI. For complete blots see **Fig. S4B**.

At 30°C in a background carrying wild-type *SAE2*, DSB steady state levels are also somewhat increased in *dot1*Δ (Fig. 4C-i), yet levels and kinetics of dHJ intermediates (Fig. 4D, C-ii) as well as crossover and non-crossover products are essentially indistinguishable from wild type (Fig. 4E, C-iii,iv) (47). Notably, in some *dot1*Δ cultures at the time of initial dHJ detection, IS-dHJs are increased over IH-dHJs, consistent with a role of *DOT1* in minimizing strand exchange between sister chromatids during early meiosis, when nucleus-wide DSB abundance is low (Fig. S4A, see t = 2.5 h in bottom panel) (6, 48).

Under meiotic heat stress, steady state DSBs are also somewhat increased in *dot1*Δ, total dHJs appear with similar timing as in the wild type, but rather than persisting, they have been resolved by t = 11 h (Fig. 4D, C-ii). Furthermore, although both IH- and IS-dHJ turn over, the relative abundance of dHJs between sister chromatids is substantially increased in *dot1*Δ compared to those between homologues at all times (Fig. 4D,H; Fig. S4A). This suggests that under heat stress, either additional IS-dHJs are formed, potentially derived from additional DSBs formed in *dot1*Δ, and/or IH-dHJs are resolved while some IS-dHJs persist. Importantly, under heat stress, crossovers are restored to substantial levels in *dot1*Δ, whereas non-crossovers remain unchanged (Fig. 4E, C-iii,iv).

All *dot1*Δ effects on DSB processing are recapitulated in the non-methylatable *h3k79r* mutant, including reduced levels of IH-dHJs (Fig. 4F,G), an increase of crossovers with minor effects on non-crossovers (Fig. 4J,I), and an increase of IS-dHJs relative to IH-dHJs, at early time points at 30°C and at all times at 34.8°C (Fig. 4F to H). By contrast, crossovers and non-crossovers at 30°C are indistinguishable from WT in *h3k79r*, consistent with effects of Dot1-mediated H3K79 methylation specifically under heat stress (Fig. 4J, I; Fig. S4B). Notably, intersister dHJs compared to interhomolog dHJs are even more abundant in *h3k79r* compared to *dot1*Δ, possibly indicating that the amino acid at position 79 affects DSB processing apart and beyond the presence of the methylation mark (Fig. 4F, H). Together, these findings indicate that under standard conditions, Dot1 via H3K79 methylation attenuates DSB formation and intersister recombination specifically during early meiosis, when DSB abundance is low (6, 48). During heat stress, Dot1-mediated H3K79 methylation further is critical for attenuating resolution of IH-dHJs into crossovers, potentially as part of its checkpoint function, even though the checkpoint effect does not depend on modulation of recombination.

## Discussion

The sensitivity of meiotic divisions to moderate heat stress is a widely conserved phenomenon across diverse organisms (18–24). Our results demonstrate that this arrest reflects an active surveillance response to defects in DSB-dependent chromosome interactions. This surveillance mechanism exhibits hallmark features of a classical checkpoint: It monitors defects associated with DSB processing, as evidenced by the restoration of meiotic divisions when recombination is genetically eliminated. Loss of checkpoint components, including *DOT1* and *DOT1*-mediated H3K79 methylation, restores meiotic divisions in a substantial fraction of cells. *DOT1* functions in checkpoint control independently of its recombination roles, since arrest bypass occurs without altering DSB processing in the resection-defective *sae2*Δ mutant. The physiological significance of this checkpoint is underscored by the severe viability defects observed when checkpoint function is compromised, indicating that the checkpoint ensures genome integrity by halting recombination and cell divisions until permissive temperatures allow proper completion of meiotic chromosome interactions.

### A chromatin-based surveillance mechanism distinct from the cytoplasmic heat shock response

Although the meiotic heat stress response shares features of both the meiotic recombination and DNA damage checkpoints (49, 50), it diverges from previously characterized surveillance mechanisms in important ways. Like the recombination checkpoint, it responds to Spo11-induced events including recombination defects (1), yet it utilizes only a subset of recombination checkpoint components. Thus, *dot1*Δ restores meiotic divisions under heat stress, while meiotic checkpoint mutant *pch2*Δ exacerbates heat sensitivity rather than suppressing it (12). Similar to the DNA damage checkpoint (49), the heat stress response requires Dot1-mediated methylation of histone H3 at lysine 79, yet it operates independently of H3K79me reader Rad9^53BP1^ (15, 31) and instead activates the meiosis-specific Rad53-paralogue Mek1, a feature shared with the meiotic recombination checkpoint (44). Importantly, the meiotic heat stress response arrests wild-type meiosis under physiologically relevant conditions, distinguishing it from the recombination checkpoint that previously was analyzed under mutation-induced conditions (10).

The meiotic heat stress response fundamentally differs from the canonical heat shock response in mitotically growing cells. In vegetative cells, temperatures above ∼37°C trigger cytoplasmic accumulation of misfolded nascent proteins sequestering Hsp70, relieving Hsf1 transcription factor inhibition, thereby triggering compensatory gene expression that enables cells to exit from transient G1 arrest and resume divisions at elevated temperatures (32, 36, 51, 52). By contrast, the meiotic heat stress response is triggered by chromatin-associated defects, likely stalled DNA recombination intermediates, that are detected in context of H3K79 methylated nucleosomes (53). This chromatin-based sensing mechanism activates a checkpoint that permanently arrests meiotic progression until temperatures return to permissive levels.

### Dual responses to heat stress during wild-type meiosis

The heat-induced meiotic arrest likely responds to recombination defects rather than those in pairing or synapsis, as *spo11-yf* bypasses the arrest despite causing substantial pairing and synapsis defects (54–57). Given that ∼90% of histone H3 molecules carry at least one methylation on K79, with ∼50% of H3 being trimethylated (58), it seems likley that stalled recombinosomes together with methylated H3K79 deform the chromatin fiber (59) thereby triggering downstream checkpoint signaling. At the same time, we cannot exclude that stalled recombinosomes promote de novo methylation of H3K79. Involvement in heat-induced meiotic arrest of Mek1 and one of its substrates, the transcription factor Ndt80 (14) further suggests that H3K79 methylation plays a direct role in sensing defects along the chromatin fiber, rather than influencing meiotic progression indirectly through transcript elongation (60).

Apart from the checkpoint-mediated arrest at the mononuclear G2/prophase stage described above, heat stress elicits a distinct, second cellular response, with a subset of cells exhibiting cellular deterioration characterized by irregular cell morphologies. The latter deteriorative response occurs independently of DSB formation and is unaffected by checkpoint status. These responses to heat stress appear to be mutually exclusive and the choice between them is potentially determined by cell cycle stage and/or cell-to-cell variability in stress responsiveness.

### Dot1 regulates recombination independently of its checkpoint role

Our findings also reveal roles of *DOT1* in meiotic recombination under both permissive and moderate heat stress conditions independent of its checkpoint function. *DOT1* attenuates DSB formation, promotes choice of the homologue as recombination partner, and suppresses crossover-specific resolution of double Holliday junctions. The latter finding provides strong evidence for the crossover-specific fate of interhomolog dHJs, a conclusion previously inferred from mutant meiosis (3, 4). Although some *dot1*Δ effects on recombination are detectable in the absence of arrest, they may still be part of the checkpoint response. Attenuation of DSB formation and dHJ resolution likely minimize genomic instability, and limiting dHJ formation between sister chromatids may prevent chromosome missegregation.

A *DOT1* role in preventing strand exchange with the sister chromatid, mediated by *RAD54*, was previously inferred genetically from its effects on meiotic progression in *dmc1*Δ (10, 47). Our physical analysis now directly demonstrates that *DOT1* minimizes intersister strand exchange during wild-type meiosis in presence of *DMC1*, likely to allow repair of extra DSBs generated in *dot1*Δ. *DOT1* is particularly critical for suppressing recombination between sister chromatids when DSB abundance is low, such as during early meiosis and under heat stress. This context-dependent regulation parallels previous observations that specialized DSB processing factors, including Tel1^ATM^, Pch2^TRIP13^, and Mph1^FANCM^, are indispensable under conditions when DSB abundance is low (6, 48).

Our finding that *DOT1* limits DSB formation contrasts with an earlier report suggesting that *DOT1* promotes DSB formation, at least in a mutant background compromised for DSB formation (47). This discrepancy is likely explained by increased intersister repair in *dot1*Δ, particularly under low DSB conditions - a function identified in the present study. Elevated DSB levels in the *dot1*Δ *sae2*Δ double mutant further argue against a model in which *DOT1* mediates compensatory DSB increases in response to defective homologue engagement, as *sae2*Δ lacks the relevant feed-back mechanism (61).

### Evolutionary conservation of the meiotic heat stress response

Several lines of evidence suggest that chromatin-based detection of recombination defects during heat stress, as discovered here, is conserved across eukaryotes. A checkpoint role during defective meiosis has been documented for *DOT1* in both yeast and metazoans (10, 62), supporting evolutionary conservation of stress-responsive meiotic surveillance mechanisms. In plants, cytological markers of DSBs are reduced or eliminated under heat stress (63, 64).

Moderate heat stress prolongs leptonema in *Allium* (23), and pachynema in *Arabidopsis*, through activation of the ATM checkpoint kinase (65). *Arabidopsis* ATM is also essential for maintaining chromosome integrity during heat-stressed meiosis (64). Notably, yeast Tel1^ATM^ shares functional similarities with Dot1, including limiting DSB formation (66) and promoting meiotic arrest in response to unresected DSBs (11). These ATM functions during meiosis are conserved in higher eukaryotes (67, 68).

### Concluding remarks

We have identified novel functions of a checkpoint that serves dual protective roles during heat-stressed wild-type meiosis: preventing premature resolution of interhomolog double Holliday junctions and blocking meiotic divisions. Our finding that the heat stress response is governed by a specific genetic program reveals that thermal sensitivity of meiosis is genetically tunable.

These findings have important implications for ensuring reproductive success and evolutionary adaptation in the context of rising global temperatures.

## Supporting information

Joshi-Supplemental Figures S1toS4

## Acknowledgments

We thank J. Mason Allen, Marta Makuszewski-Lavery, and Robin E. Woods for assistance with preliminary experiments, Akira Shinohara, Nancy Kleckner, and Rima Sandhu for strains, Aaron Severson, Judith Yanowitz, Anton Komar, Alan Tartakoff, and Börner lab members for discussion.

