## Supplementary material for "An H3K79 Methylation-Dependent Checkpoint Blocks Holliday Junction Resolution and Meiotic Divisions under Heat Stress": Joshi-Supplemental Figures S1toS4

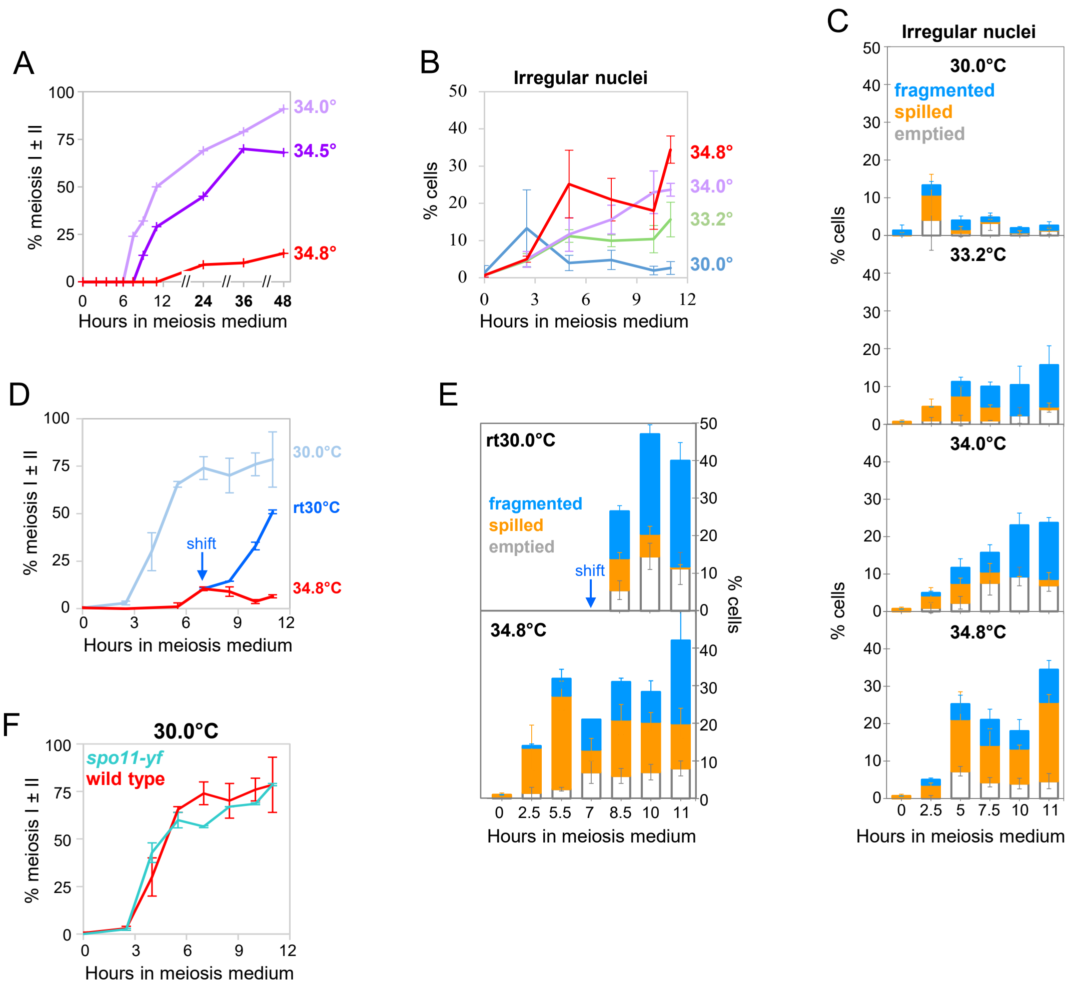


**Fig. S1. Meiotic cell divisions and irregular morphologies of undivided cells at different temperatures.**

(A) Moderate heat stress (34.8°C) triggers robust meiotic arrest for up to 48 hours, while subcultures derived from the same master culture complete meiosis efficiently at slightly lower temperatures.

(B) Kinetics of appearance of irregular nuclear morphologies at temperatures ranging from 30°C to 34.8°C. n = 3; error bars are SDs. See **Fig. S1C** for individual nucleus classes.

(C) Classes of undivided meiotic cells with irregular nuclear morphologies at temperatures ranging from 30.0°C to 34.8°C. Cultures are the same as in **Fig. 1C** (n = 3; error bars are SDs. For representative images see **Fig. 1A, iii-v**).

(D) Meiotic progression of wild type returned to permissive conditions following transient exposure to moderate heat stress (34.8°C) between t = 2 h and 7 h. n = 2; error bars indicate range.

(E) Classes of undivided meiotic cells with irregular nuclear morphologies following shift of an aliquot from 34.8°C to 30.0° (top) and at 34.8°C (bottom). n = 2; error bars indicate range.

(F) Meiotic cell divisions in catalytically dead *spo11-yf* and wild type at 30°C. For progression at 34.8°C of cultures derived from the same *spo11-yf* and WT master cultures see **Fig. 1H**. n = 2; error bars indicate range.

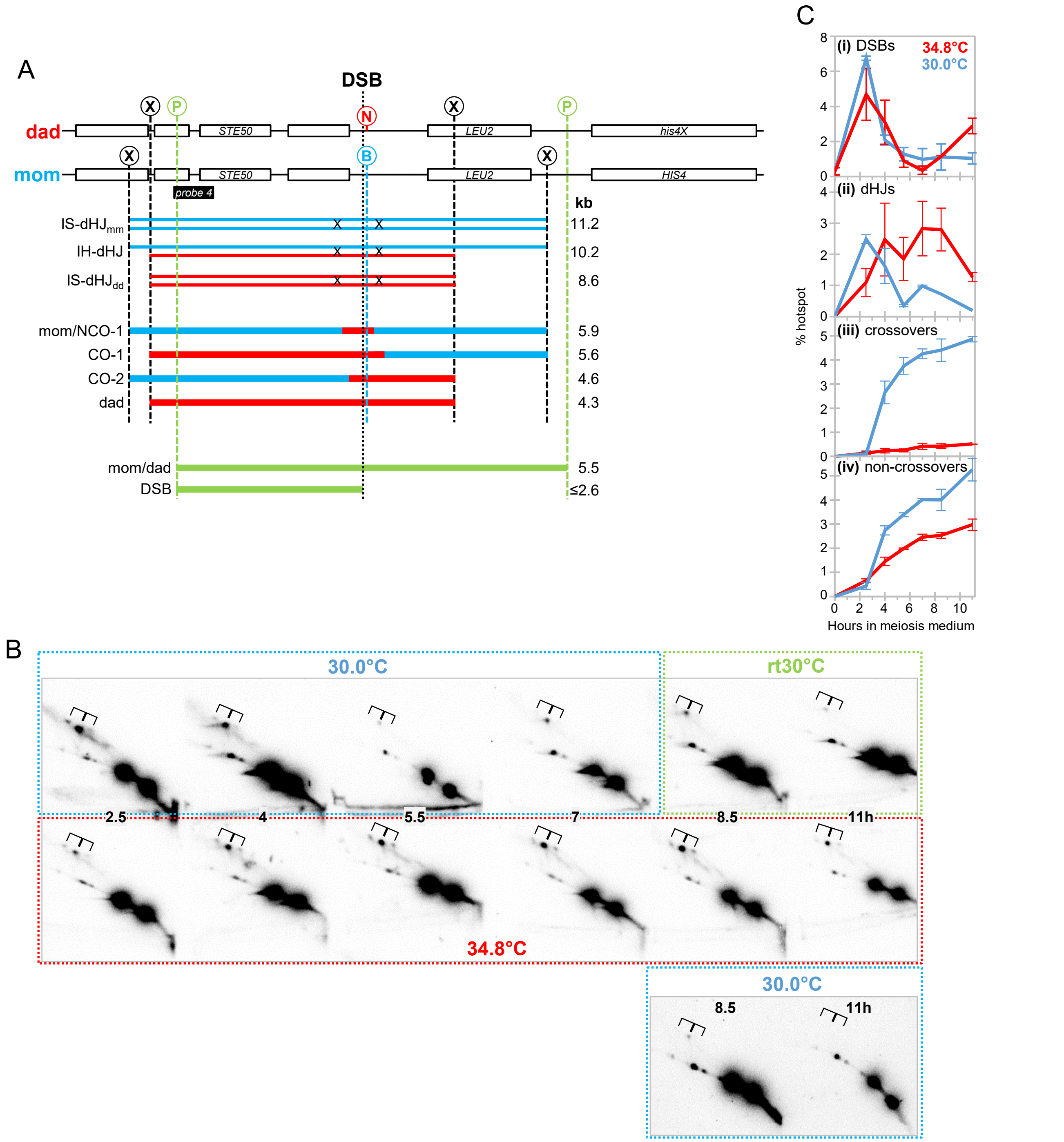


**Fig. S2. Effects of heat stress on recombination.**

(A) Diagram of the *HIS4LEU2* recombination hotspot. "mom" and "dad" refer to the two DSB hotspot versions in the heterozygous parental strain that differ in the positions of flanking XhoI sites (circled X), as well as a restriction site polymorphism adjacent to the central DSB site where "dad" carries an NgoMIV site (circled N) and "mom" a BamHI site (circled B). The BamHI polymorphism was used to detect mom fragments that had undergone gene conversions into the BamHI-refractory NCO-1 product. "Probe 4" was used for all Southern blot analyses (38). Recombination species in blue and red indicate dHJs between homologues or sister chromatids [with Holliday junctions symbolized by a black X] (top), parental restriction fragments as well as crossover and non-crossover products (middle). Green fragments indicate parental band and DSBs as assayed by PvuII digest (circled P).

(B) Representative 2D gel Southern blot analyses of recombination intermediates at the *HIS4LEU2* hotspot between 2.5 h and 11 h in a wild-type strain undergoing meiosis at 30°C, 34.8°C, or returned to 30°C after 5 hours at 34.8°C (rt30°C). Dotted frames in blue (30°C), red (34.8°C), and green (rt30°C) symbolize the respective incubation conditions.

(C) Quantitative analysis of recombination intermediates and products at the *HIS4LEU2* hotspot at 34.8°C and 30°C. Data are from **Fig. 2D** without axis adjustment.

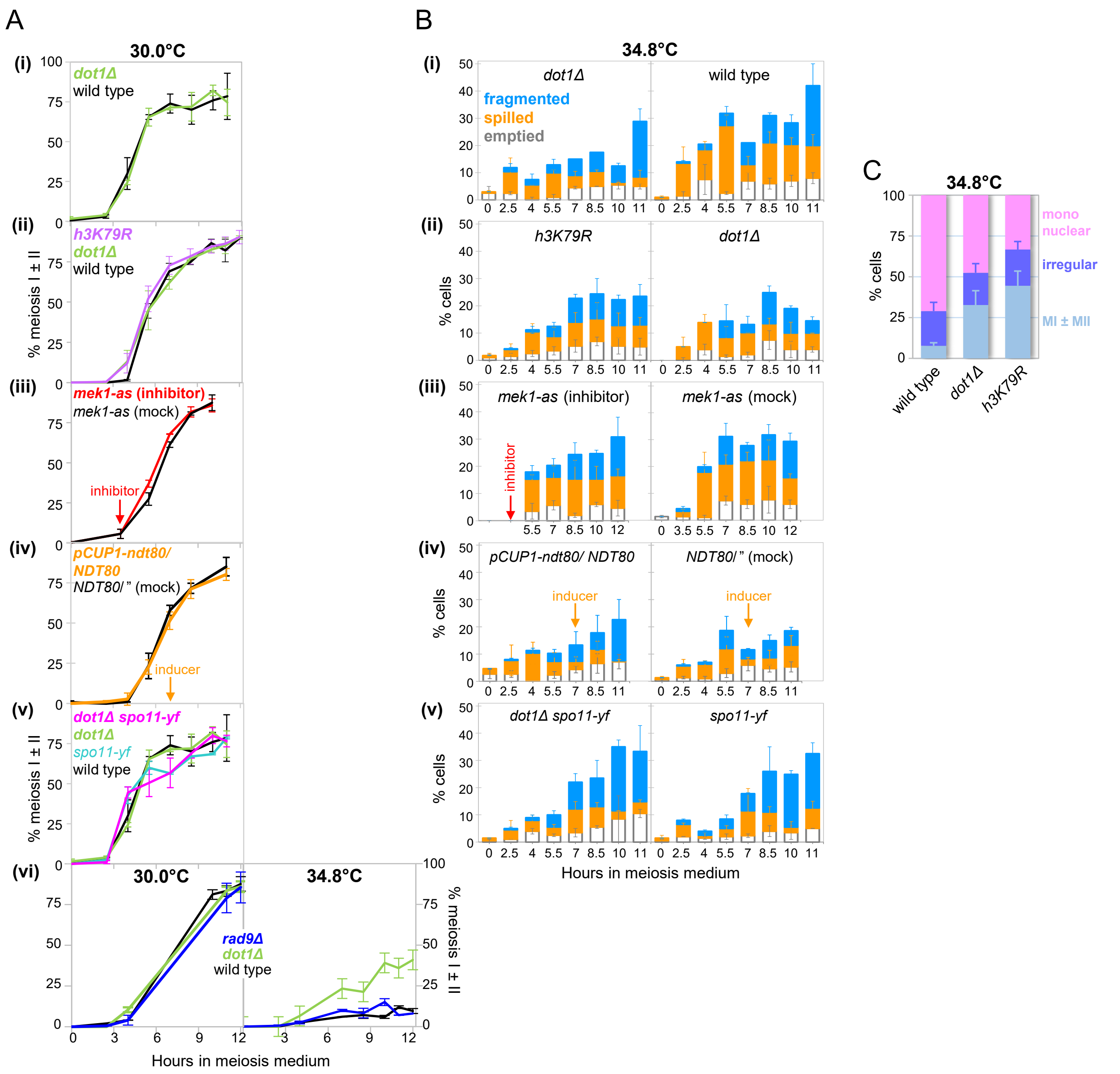


**Fig. S3: Mutant effects on meiotic progression and nuclear morphologies under heat stress and standard conditions.**

(A) Average percentage of cells undergoing one or both meiotic divisions under standard conditions (30°C) in cultures of the indicated genotypes as determined by epifluorescence microscopy of DAPI-stained, fixated cells. **(i)** *dot1Δ* versus WT (n = 2), **(ii)** *h3k79r* (n = 4) versus *dot1Δ* and WT (n = 2), two *h3k79r* strains (NOJ1547, NOJ1548) were each analyzed in duplicate, with essentially identical results, **(iii)** ATP analog-sensitive *mek1-as1* plus inhibitor added at t = 3.5 h versus *mek1-as1* mock-treated subculture (n = 3), **(iv)** culture heterozygous for *pCUP1-ndt80*/*pNDT80-NDT80* induced at t = 7 h with copper sulfate compared to an identically-treated homozygous *pNDT80-NDT80/"* (WT) culture (n = 3), **(v)** *spo11-yf dot1Δ* double mutant compared to the respective single mutants and wild type (n = 2). **(vi)** *rad9Δ* versus *dot1Δ* and WT (n = 2) at 30°C (left) and 34.8°C (right). Error bars indicate range or SD, respectively, for n = 2 and n ≥ 3. For the corresponding cultures at 34.8°C see **Fig. 3B**.

(B) Average frequencies of meiotic cells with irregular nuclear morphologies at 34.8°C, in cultures of the indicated genotypes. For numbers of cultures analyzed in each experiment, see legend **Fig. 3B**. Wild-type data in **Fig. S3B-i** are from the same experiment at 34.8°C as shown in **Fig. S1E**. Error bars indicate range or SD, respectively, for n = 2 and n ≥ 3.

(C) Meiotic arrest bypass (MI ± MII) at 34.8°C in *h3k79r* and *dot1Δ* reduces the abundance of mononucleate cells but leaves irregular morphologies unchanged. Percentages of regular (mononuclear or MI±MII) or irregular cell morphologies are averages of t = 10 h and 11 h. Cultures are the same as in **Fig. 3B-ii** i.e., *h3k79r* (n = 4) versus *dot1Δ* and WT (n = 2). Error bars are SD.

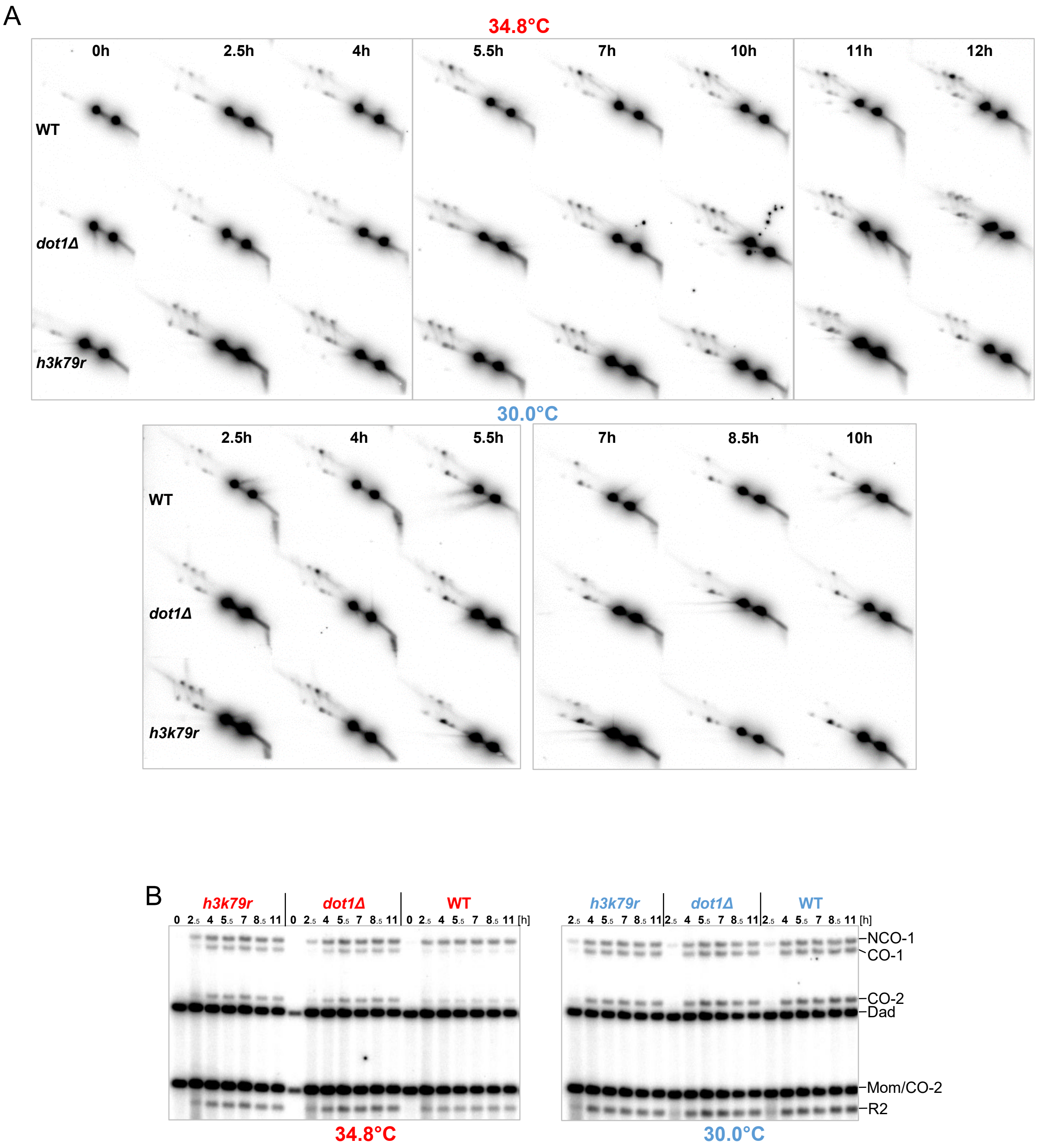


**Fig. S4. Role of histone H3K79 methyltransferase Dot1 in recombination.**

(A) Complete 2D gel Southern blots of WT, *dot1Δ*, and *h3k79r* at 34.8°C (top) and 30°C (bottom).

(B) Complete 1D gel Southern blots of genomic DNA digested with XhoI and BamHI at 34.8°C (left) and 30.0°C (right). For relevant fragments see diagram in **Fig. S2A**.
